# Young adult microglial deletion of C1q reduces engulfment of synapses and prevents cognitive impairment in an aggressive Alzheimer’s disease mouse model

**DOI:** 10.1101/2025.09.13.676049

**Authors:** Tiffany J. Petrisko, Shu-Hui Chu, Angela Gomez-Arboledas, Blossom Zhang, Andrea J. Tenner

**Affiliations:** Department of Molecular Biology & Biochemistry, University of California, Irvine, Irvine, CA; Department of Neurobiology and Behavior, University of California Irvine, Irvine CA, USA; Department of Pathology and Laboratory Medicine, University of California, Irvine, School of Medicine, Irvine, CA, USA

**Author notes:** **Corresponding Authors:** Andrea J. Tenner, University of California, Irvine, 3205 McGaugh Hall, Irvine, CA 92697-3900, Tiffany J. Petrisko, University of California, Irvine 3205 McGaugh Hallm Irvine, CA 92697-3900. **Abbreviations**: Aβ (amyloid beta), AD (Alzheimer’s disease), APP (amyloid precursor protein), Arc (Arctic48), AU (Airy Unit), CLU (Clusterin), C1q (Complement component 1q), CR1 (Complement receptor 1), C3 (Complement component 3), C5aR1 (complement component C5a receptor 1), DAMP (Damage Associated Molecular Patterns), fAβ (fibrillar amyloid beta), GFAP (Glial Fibrillary Acidic Protein), GWAS (Genome-wide association studies), Iba1 (Ionized Calcium-binding adapter molecular 1), Immunofluorescent (IF), Megf10 (Multiple EGF-like-domains 10, MerTK (Mer tyrosine kinase), NfL (Neurofilament Light), OLM (Object Location Memory), PtdSer Phosphatidylserine), ThioS (Thioflavin S), TREM2 (Triggering Receptor Expressed on Myeloid Cells 2), Vglut1 (vesicular glutamate transporter 1).

**Keywords:** Alzheimer’s disease, complement, C1q, microglia, synapse, cognition, amyloid

## Abstract

C1q is a multifunctional protein, including its role as the initiating protein of the classical complement cascade. While classical pathway activation is involved in synaptic pruning during development of the nervous system, it also contributes to enhanced inflammation and cognitive decline in Alzheimer’s disease (AD). Constitutive genetic C1q deficiency has been shown to reduce glial activation and attenuate neuronal loss in AD mouse models, but the specific contributions of microglial C1q to AD pathology while avoiding deficits during post-natal development remain to be determined. To dissect specific role(s) of microglial C1q in AD progression, we crossed the Cx3cr1^CreERT2^ mouse model that deletes C1q from microglia in young adulthood (8 weeks of age) to the aggressive Arctic48 (Arc) amyloidosis mouse model. At 10 months, young adult microglial C1q deletion (Arc C1q^ΔMG^) rescued cognitive deficits in spatial memory, despite unchanged amyloid plaque burden. Furthermore, Arc C1q^ΔMG^ mice exhibited reduced hippocampal C3 protein levels without altering C3 mRNA. No changes were observed in C5aR1, astrocyte GFAP, or microglial Iba1 protein expression. However, Arc C1q^ΔMG^ mice demonstrated region specific reductions in microglial synaptic engulfment, alongside decreased phagolysosome-associated amyloid in both microglia and astrocytes, and reduced compaction of amyloid within the hippocampus. These findings support a role for C1q in astrocytic C3 induction and the engulfment of both synapses and amyloid. Importantly, young adult microglial C1q inhibition confers cognitive benefits without exacerbating amyloid pathology, suggesting a therapeutic window in which targeting microglial C1q may mitigate neuroinflammation and synaptic loss during the later stages of AD.

**Main Points:**

- Young adult deletion of microglial C1q reduced engulfment of Vglut1+ synapses and preserved spatial cognition at 10 months of age in AD mice.
- Fibrillar amyloid plaques nor soluble or insoluble Aβ levels in the hippocampus were affected by young adult microglial loss of C1q despite reduced phagocytosis of amyloid by both microglia and astrocytes.
- Loss of microglial C1q reduced C3 levels but did not affect astrocytic GFAP expression in AD mice indicating altered polarization of astrocytes.

## Introduction

Alzheimer’s disease (AD) is the most common form of dementia in the elderly. In the United States alone, over 7.2 million individuals suffer from AD, with the affected number anticipated to reach 13.8 million by 2060 (Alzheimer’s Association, 2025). Neuropathologically, AD is characterized by the extracellular accumulation of amyloid-beta (Aβ) in amyloid plaques and the intracellular accumulation of hyperphosphorylated tau, known as neurofibrillary tangles, as well as synaptic loss (DeTure and Dickson, 2019).

Emerging evidence has highlighted the critical role of neuroinflammation and innate immune activation in AD progression (Kinney et al., 2018; Heneka et al., 2025). The complement system, an essential component of the innate immune response, has gained significant attention (Negro-Demontel et al., 2024; Nimmo et al., 2024; Tenner and Petrisko, 2025). Genome-wide association studies (GWAS) have repeatedly linked several complement components to AD susceptibility, including *CLU* (clusterin) and CR1 (Lambert et al., 2009), and many complement factors colocalize with fAβ plaques in both human and mouse brain tissue (reviewed in Schartz and Tenner (2020)).

Activation of the classical complement pathway begins via engagement of the macromolecular complex C1, comprising the recognition component, C1q, associated with the serine proteases, C1r and C1s. C1 can be activated by C1q binding to the Fc region of IgG or IgM immunoglobulin bound to an antigen or by damage- or pathogen-associated molecular patterns, apoptotic cells, DNA, or damaged mitochondria. Furthermore, complement activation occurs in response to the pathological hallmarks of AD, namely fibrillar Aβ (fAβ) (Afagh et al., 1996; Velazquez et al., 1997; Tacnet-Delorme et al., 2001) and hyperphosphorylated tau (Shen et al., 2001).

Classical pathway activation results in the generation of C3a, C3b and subsequently terminal pathway products C5a and C5b-9. Previous studies have demonstrated a role for C1q and complement activation in removal of excess synapses during development (Stevens et al., 2007; Chu et al., 2010; Schafer et al., 2012) and in synaptic plasticity in the adult homeostatic brain (Wang et al., 2020), a process defined as synaptic pruning. C1q expression increases with age, injury, and disease (Stephan et al., 2013; Tenner and Petrisko, 2025), and activation of C1 and C3 have also been implicated in AD as a mechanism of synaptic loss and cognitive impairment (Hong et al., 2016; Wu et al., 2019), with constitutive knockouts of C1q restoring synaptic density and reducing pathology in both tauopathy (Dejanovic et al., 2022) and amyloidosis mouse models (Fonseca et al., 2004).

C1q itself is composed of six heterotrimers, each consisting of A, B, and C polypeptide chains, all of which are necessary for proper assembly and function of C1q and contain multiple recognition sites (Sontheimer et al., 2005; Roumenina et al., 2006; Bally et al., 2013). Within the CNS, C1q is produced almost exclusively by microglia (Fonseca et al., 2017; Scott-Hewitt et al., 2024); however, the contribution of microglial C1q to the progression of AD remains unknown and the use of constitutive knockout models have limited interpretation due to the known developmental and homeostatic roles of C1q. Here, we examined the long-term effects of microglial C1q deletion during young (8 weeks) adulthood in the aggressive Arctic AD mouse model using the Cx3cr1^CreERT2^ knock in mouse, which deletes C1q from microglia by 8 weeks of age independent of tamoxifen treatment (Fonseca et al., 2017). Our results show that deletion of microglial C1q in young adulthood reduces engulfment of hippocampal synapses and attenuates cognitive deficits in 10-month-old Arctic AD mice. Furthermore, we demonstrate that while C1q deletion fails to reduce amyloid load at 10 months of age, the structure of amyloid plaques and phagocytosis of amyloid by glia is altered.

## Methods

### Animals

All animal experimental procedures were approved by the Institutional Animal Care and Use Committee of University of California, Irvine, and performed in accordance with the NIH Guide for the Care and Use of Laboratory Animals.

All mice were housed under a 12-h light/dark cycle with ad libitum access to food and water. Arctic48 (Arc) mice carry the human APP transgene with the Indiana (V717F), Swedish (K670N/M671L), and Arctic (E22G) mutations (under the control of the platelet-derived growth factor-ß promoter) on the C57BL/6 background, resulting in the production of amyloid plaques as early as 2-3 months of age (Cheng et al., 2004). To generate microglial specific knockouts of C1q, WT *C1qa^FL/FL^* and Arc *C1qa^FL/FL^* mice were crossed to B6.129P2(Cg)- *Cx3cr1^tm2.1(cre/ERT2)Litt^*/WganJ animals, (Jackson, stock #021160) to generate WT *C1qa^FL/FL^* or Arc C1qa^FL/FL^ *Cx3cr^CreERT2^* mice (abbreviated C1q^ΔMG^), as previously described (Fonseca et al., 2017). Microglial deletion of C1q in Cx3cr1^CreERT2^ mice was previously shown to occur independently of tamoxifen treatment by 2 months of age (Fonseca et al., 2017). Arctic48 mice were originally obtained from Dr Lennart Mucked (Gladstone Institute, San Francisco, CA, USA). Both males and females were used in all experiments. WT *C1qa^FL/FL^* and Arc *C1qa^FL/FL^*mice, with and without Cx3cr1^CreERT2^ were housed together by sex.

### Blood Collection and Plasma Isolation

Blood samples were collected at two time points: 7 months and 10 months of age. At 7 months, blood was obtained via submandibular vein puncture using a 5mm lancet to collect ∼ 100uL of blood. Pressure was immediately applied to stop bleeding. At 10 months, blood was collected immediately prior to perfusion via cardiac puncture using a 25G syringe. In both cases, blood was collected into EDTA-containing Eppendorf tubes (10mM final concentration) maintained on ice. Blood was centrifuged at 2348 g for 10 minutes at 4°C to separate the plasma. The plasma was immediately aliquoted and stored at -80°C until further analysis.

### Tissue Collection

Mice were deeply anesthetized with isoflurane and transcardially perfused with cold phosphate buffered saline (PBS). Brains were rapidly harvested and were split along the midline to allow for comparison of the same animal across various biochemical assays. For qPCR, western blot, or Aβ MSD analysis, the hippocampus and cortex were isolated, immediately placed on dry ice, and stored at -80°C. For immunofluorescent analysis, half brains were post-fixed for 24 hours in 4% paraformaldehyde/PBS at 4°C and transferred to 0.02% sodium azide/PBS for long-term storage at 4°C.

### Immunofluorescence (IF)

Brains were cut into 30μm thick coronal sections on a Leica VT1000S vibratome and stored in 0.02% sodium azide/PBS at 4°C for long-term storage. Free-floating sections were washed in 1X PBS. When staining for 6E10, sections underwent antigen retrieval in 50mM citrate buffer, pH 6.0 for 20 minutes in 80°C water bath while C5aR1 sections underwent antigen retrieval in 50mM TBS, 0.05% Tween, pH 9.0 in 80°C water bath for 30 min and allowed to cool for 10 minutes at room temperature (RT). Sections were then incubated in blocking buffer for 1 hour at RT shaking, followed by incubation with primary antibodies in blocking buffer at 4°C overnight (**Table 1**). Blocking buffer consisted of 5% normal goat serum (NGS), 2% bovine serum albumin (BSA,) 0.1% Triton in PBS, except for C1q staining, which used 2% BSA, 0.1% Triton in PBS. Sections were then washed in 1X PBS before secondary antibody incubation (1:500, **Table 1**) for 1 hour at RT, shaking. Sections were then washed in 1XPBS and mounted onto a slide and coverslipped using Liquid Antifade Mounting Medium (Vectashield, Vector) and sealed with nail polish. To visualize plaques, tissues were incubated for 10 min with Thioflavin S (0.5% in MilliQ water, Sigma-Aldrich #T1892) or AmyloGlo (1:100, Biosensis #TR-300-AG in PBS). Within each experiment, 1-3 sections/mouse were stained and processed at the same time, using batch solutions.

**Table 1:** Antibodies for Immunofluorescence and Western Blotting.

| **Antibody/ Stain** | **Host Species** | **Dilution** | **Antigen Retrieval** | **Supplier-Catalog #** | **Catalog #** | **Blocking Buffer** |
| --- | --- | --- | --- | --- | --- | --- |
| **B-actin** | Mouse | 1:7000 |  | Sigma | A1978 | 3% milk |
| **C1q clone 27.1** | Rabbit | Hybridoma supernatant |  | Tenner Lab |  | 2% BSA/ 0.1% Triton X-100 in 0.1M PBS |
| **C1q clone 1151** | Rabbit | 5.1µg/mL |  | Tenner Lab |  | 5% milk |
| **C3 clone 11H9** | Rat | 1:50 |  | Hycult | HM1045 | 5% NGS/ 2% BSA/<br>0.1% Triton X-100 in<br>0.1M PBS |
| **C5aR1 (CD88 10/92)** | Rat | 1:1000 | 50mM TBS,<br>0.05% Tween,<br>pH 9.0 | BioRad | MCA2456 |  |
| **GFAP** | Rabbit | 1:2900 |  | Dako | Z0334 |  |
| **Iba1** | Rabbit | 1:1000 |  | Wako | 019-1974 |  |
| **CD68 (clone FA-11)** | Rat | 1:1000 |  | BioRad | MCA1957 |  |
| **Lamp2** | Rat | 1:1000 |  | Abcam | Ab13524 |  |
| **6E10** | Mouse | 1:1000 | 50mM citrate buffer, pH 6.0 | Biolegend | 803001 |  |
| **Vglut1** | Guinea Pig | 1:1000 |  | Millipore | AB5905 |  |
| **Thioflavin S** |  | 0.50% |  | Sigma | T1892 |  |
| **AmyloGlo** |  | 1:100 |  | Biosensis | TR-300-AG |  |
| **Alexa Fluor IgG (H+L) Goat α Guinea Pig 488** | Goat | 1:500 |  | Invitrogen | A11073 |  |
| **Alexa Fluor IgG (H+L) Goat α Mouse 488** | Goat | 1:500 |  | Invitrogen | A11029 |  |
| **Alexa Fluor IgG (H+L) Goat α Rat 555** | Goat | 1:500 |  | Invitrogen | A21434 |  |
| **Alexa Fluor IgG (H+L) Goat α Rabbit 555** | Goat | 1:500 |  | Invitrogen | AA21428 |  |
| **Alexa Fluor IgG (H+L) Goat α Rabbit 647** | Goat | 1:5000 |  | Invitrogen | A21244 |  |
| **Peroxidase AffiniPure® F(ab')<sub>2</sub> Fragment Donkey Anti-Rabbit IgG (H+L)** | Donkey | 1:5000 |  | Jackson Labs | 711-036-152 | 3% or 5% milk |
| **Peroxidase AffiniPure® F(ab')<sub>2</sub> Fragment Donkey Anti-Mouse IgG (H+L)** | Donkey | 1:5000 |  | Jackson Labs | 715-036-150 |  |

### Imaging and Quantification

To confirm microglial deletion of C1q within the brain, images of the dentate gyrus were obtained at 10x with the Zeiss Axiovert 200 inverted microscope (Zeiss) and images acquired with a Zeiss Axiocam high-resolution digital camera (1300 x 1030-pixel resolution) using ZEN 2.6 software. The intensity of C1q staining was quantified in 4-6 mice/genotype using ImageJ. Five regions of interest (ROI) squares were defined randomly within the molecular layer of the dentate gyrus and the mean pixel intensity per ROI was determined. The mean C1q intensity of each animal was obtained by averaging the intensities of the 5 ROIs and averaged across 1-2 sections/mouse.

Low magnification images of Thioflavin S staining were obtained using the Zeiss Axio Scan.Z1 slide scanner using a 10x objective. Confocal hippocampal tile scan z-stacks of triple labeled C5aR1/Iba1/Amylo-Glo and C3/GFAP/Amylo-Glo were captured using a Leica Sp8 confocal microscope equipped with a 20 x 0.8 numerical aperture (NA) objective using a 1μm z-step with a 1X zoom. All confocal imaging with the Leica Sp8 microscope utilized a pixel resolution of 1024 x 1024 and pinhole set to 1 Airy Unit (AU) and all settings were maintained within an imaging set for consistency. Using Imaris 9.7 (Oxford Instruments), masks were made of the hippocampus and surfaces were rendered for each channel and the percentage of hippocampal surface area or volume per channel was calculated. Three hippocampal sections (dorsal, middle, and ventral) were imaged from each mouse and averaged together, with 4-8 mice per genotype.

Images for the engulfment of excitatory synapses (Vglut1) by microglial were obtained with a 63x 1.4 NA objective with a 0.3μm step size using a 3.5x zoom. Using Imaris 10.0, synaptic engulfment was quantified in 12 microglia cells/ mouse per region (CA1 or CA3) with 4-7 mice per genotype. Lysosome associated synaptic engulfment was defined as the co-localization (≤200 nm distance) of Vglut1 + puncta (detected using spots) with CD68 and Iba1 surfaces and normalized to the total volume of the image.

Microglial (Iba1/CD68/6E10) and astrocytic (GFAP/Lamp2/6E10) phagocytosis of amyloid were acquired using a 40 x 1.3 NA objective, with a 0.5μm step size and a 3.0 zoom. In Imaris 10.0, surfaces were rendered for each channel, and the phagocytosis index was defined as the co-localization of Iba1, CD68, and 6E10 (microglia) or GFAP, Lamp2, and GFAP (astrocytes) surfaces, normalized to the total volume of Aβ per image. Microglial and astrocytic engagement with amyloid plaques was also quantified as the ratio of co-localization of Iba1/6E10 or GFAP/6E10 surfaces to total Aβ within field of view. To quantify the amount of Iba1, CD68, GFAP, or Lamp2 expression surrounding plaques, the volume of each surface was normalized to the total image volume. A total of 20 randomly selected hippocampal amyloid plaques per mouse were imaged, with 4-6 mice per genotype.

### Western Blot

Hippocampi were pulverized into powder and stored at -80°C. Hippocampi were then solubilized by homogenization in 10 volumes of Tris-buffered saline (TBS, pH 7.4) containing protease inhibitor cocktail solution (Complete mini, Roche), PhosSTOP (Roche), 1mM EDTA and 1% sodium dodecyl sulfate (SDS) using a motor pestle for 5 seconds twice on ice and centrifuged at 18,400 g for 30 minutes at 4°C. Protein concentration of the supernatant was determined using the BCA protein assay (Pierce, Rockford, IL, USA). 30 μg of hippocampal protein was loaded per lane as previously described (Fonseca et al., 2017). Briefly, samples were subjected to a 10% SDS-polyacrylamide gel electrophoresis under reducing conditions. Gels were transferred at 4 °C in transfer butter (Tris-glycine, SDS, and 10% methanol) onto a polyvinylidene difluoride (PVDF, Immobilon-P, Millipore) membrane at 300 mA for 2 hours. The membrane was then blocked for 1 hour at RT in 5% nonfat dry milk in TBS-Tween (TBST), before being incubated with rabbit anti-mouse C1q (1151) (Huang et al., 1999; Fonseca et al., 2017) in 5% milk overnight at 4°C or β-actin (Sigma) in 3% milk for 2 hours at RT. After 3 washes in TBST for 5 minutes, membranes were incubated with HRP-conjugated secondary antibodies diluted to 1:5000 (Jackson Labs) for 1 hour at RT. The blots were developed using ECL2 (Pierce) and imaged using a BioRad ChemiDoc image system and quantified using Image J software (Khoury et al., 2010).

### qPCR

Total RNA from pulverized mouse brain (10-15 mg) was extracted using the RNeasy Plus Mini Kit (Qiagen). cDNA synthesis was performed using SuperScript III reverse transcriptase (Life Technologies) according to the manufacturer’s protocol. Quantitative RT-PCR was performed using the CFX Duet Real-Time PCR System and the CFX Maestro software (Bio-Rad) with the maxima SYBR/Green Master Mix (Thermo Fisher Scientific). The mouse primer sequences for *C3, C4*, *C5ar1* and *Hprt* were obtained from their corresponding reference obtained from Eurofins (Fisher Scientific) (**Table 2)**. cDNA from each hippocampus was tested in triplicate. Within each sample, ΔCt was calculated by subtracting the average of the triplicate cycle threshold (Ct) value of target gene to the average of the triplicate Hprt Ct value. Then the relative expression was calculated by fold difference (2^−ΔCt^) multiplied by 1000.

**Table 2:**
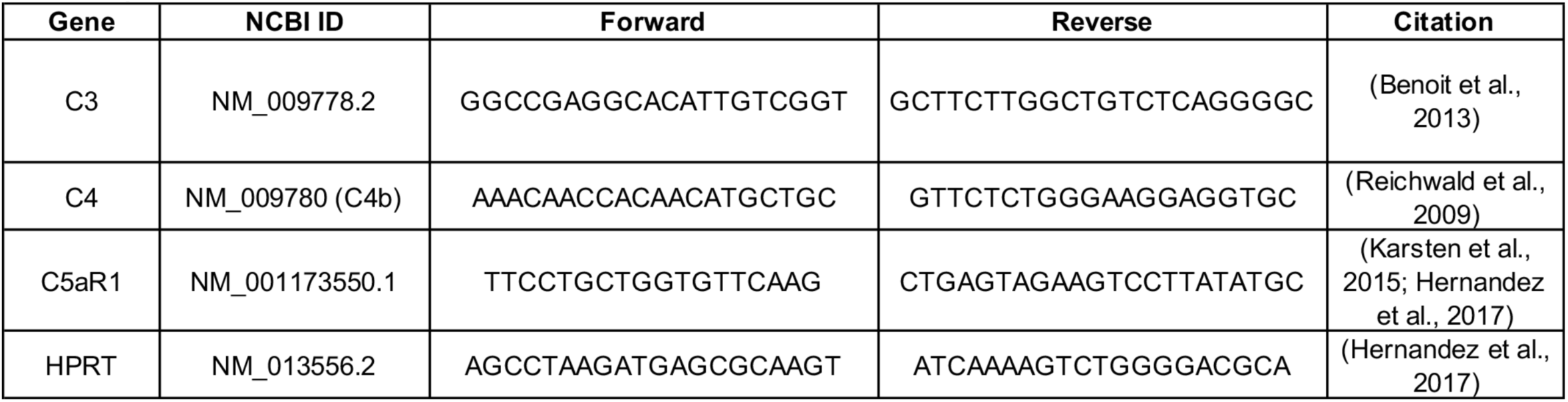
qPCR Primers.

### NfL MSD

Plasma NfL was quantified using the MesoScale Discovery R-Plex Human Neurofilament-L Assay (cat #: K1517XR-2) according to manufactures directions. Plasma was diluted 1:1 in Diluent 12. All analyzed samples were within the detection range of the assay and data is expressed in pg/mL plasma.

### Object Location Memory

At 38 weeks of age, Cx3cr1^CreERT2^ mice underwent cognitive behavioral testing. Object location memory (OLM) was conducted as previously reported (Hernandez et al., 2017). Briefly, mice were handled for 2-5 minutes per day for 4 days in the testing room and habituated to the arena (37x30x23cm), one wall marked down the center with tape as cue) covered with ∼ 1cm of sawdust bedding in dim lighting) for 5 minutes a day for 4 days, beginning on the third day of handling. The first day of habituation was recorded as an open-field trial to measure anxiety and locomotion. During the OLM training sessions, mice were placed into an empty arena for 1 minute, placed into individual cages, and 2 identical objects (large blue Legos) were placed across from one another, near the upper wall of the arenas while allowing the mice enough space to pass around the object. Mice were then reintroduced into the arena and allowed to freely explore for 10 minutes. Mice underwent identical training (objects remaining in same positions) 24 hours later. 24 hours after this final training sessions, one object was moved to the opposite corner of the arena (near side wall) to test for object location memory during a 5-minute testing period. After each session, bedding was stirred, and OLM objects were cleaned with 10% ethanol. Different bedding was utilized for males and females, and objects and arenas were cleaned with 70% ethanol and allowed to air dry when switching sexes. All mice were acclimated to room and lighting conditions for a minimum of 1 hour each day. Females were always run prior to males. During testing, OLM objects were counterbalanced to control for side preference. All trials were recorded by a mounted camera from above and interactions with objects were scored manually by blinded scorers. The percent discrimination index (%DI) was calculated [((time spent with object moved - time spent with unmoved object)/ time spent with both objects) X 100]. Mice were removed from analysis if they spent less than 7s total with the objects during training, less than 2s during testing, if they exhibited a preference for one object location over the other during training (±20 %DI), or if the performance of the mice was ± 2 standard deviations from the mean. Distance travelled, velocity, and time spent in center of arena were analyzed for all habituation sessions using EthoVision V14 (Noldus).

### Isolation of Soluble and Insoluble Aβ Fractions

To obtain the soluble and insoluble fractions of Aβ, flash frozen hippocampi were pulverized and 20-30mg of powder was homogenized in 154 μL of Tissue Protein Extraction Reagent (TPER) with 1 pellet of cOmplete^TM^, mini, EDTA-free Protease Inhibitor Cocktail (Sigma, cat#: 11836170001) and 100 μL Halt^TM^ Phosphatase Inhibitor Cocktail (Thermo Fisher, cat #: 78426) per 10mL of TPER. Samples were then centrifuged at 100,000 g for 1 hour at 4°C. The supernatant was collected as the soluble Aβ fraction and stored at -80°C. To generate the insoluble fractions, pellets from the TPER-soluble fractions were homogenized in 75uL of 70% formic acid. Afterwards, the samples were again centrifuged at 100,000 g for 1 hour at 4°C and the supernatant collected as the “insoluble” fraction and stored at -80°C. Formic acid was neutralized with neuralization buffer prior to running MSD (Butler et al., 2024).

### Aβ MSD

Hippocampal soluble and insoluble amyloid were quantified using the V-Plex Aβ Peptide Panel 1 (6E10) kit from MesoScale Discovery (cat: # K15200E) and ran according to manufacturer instructions. Soluble fractions were diluted 1:2 with Diluent 35 while insoluble fractions were diluted to a final concentration of 1:20,000 with Diluent 35 (Butler et al., 2024). Each sample was run in duplicate and normalized to the tissue weight. All samples analyzed were within the detection range of the assay and data are expressed in pg/mg tissue.

### Statistical Analysis

All statistical analyses were done with Prism V9.3 (GraphPad). Student’s t-test was utilized to compare two groups, while one-way analysis of variance (ANOVA) followed by Tukey’s post-hoc test comparing all groups was used to compare WT and Arc genotypes with or without C1q deletion. To compare plasma NfL at 7m and 10m of age, a two-way ANOVA with post hoc analysis comparing within age (simple effects analysis) was used.

## Results

### Young adult C1q microglial deletion rescues cognitive impairment in Arctic mice

WT C1qa^FL/FL^ and Arc C1qa^FL/FL^ mice bred to contain Cx3cr1^CreERT2^ (abbreviated WT C1q^ΔMG^ or Arc C1q^ΔMG^) undergo tamoxifen-independent deletion of C1q at ∼8 weeks of age (Fonseca et al., 2017). Microglial deletion of C1q was confirmed in the brain of 10m old wild type and Arc mice by immunohistochemistry and western blot analysis of the hippocampus (**Fig S1**), consistent with a previous report by Fonseca et al. (2017), which also demonstrated that C1q expression in plasma was not reduced. In order to assess the functional long-term consequences of young adult microglial C1q deletion in AD, we assessed behavioral and cognitive performance at 10 months of age. Arc mice displayed a 19% decrease in total distanced moved during five-minute open field testing compared to WT mice (p<0.05; **Fig 1A**), which was rescued in Arc C1q^ΔMG^ mice (p<0.05 between Arc-Arc C1q^ΔMG^ mice). However, differences in total distance traveled between genotypes were eliminated beginning on Day 2 of habitation (data not shown). Additionally, no differences in anxiety were noted between groups (**Fig 1B**), as assessed by the percentage of time spent in the center of the arenas during open field testing. To assess long term hippocampal-dependent spatial memory, we employed the object location memory (OLM) test (Barker and Warburton, 2011; Vogel-Ciernia and Wood, 2014; Chao et al., 2022). 24 hours after a two-day training period, WT and WT C1q^ΔMG^ animals displayed appropriate preference for the object moved to the novel location. Arc mice had a significant impairment in spatial memory compared to WT mice (p<0.01; **Fig 1C**), which was prevented in Arc C1q^ΔMG^ animals (p=0.057). No differences in distance travelled or total object interaction time during training between genotypes was observed (data not shown), demonstrating that there is long-term cognitive benefit to microglial C1q deletion during young adulthood in AD mice.

**Figure 1:**
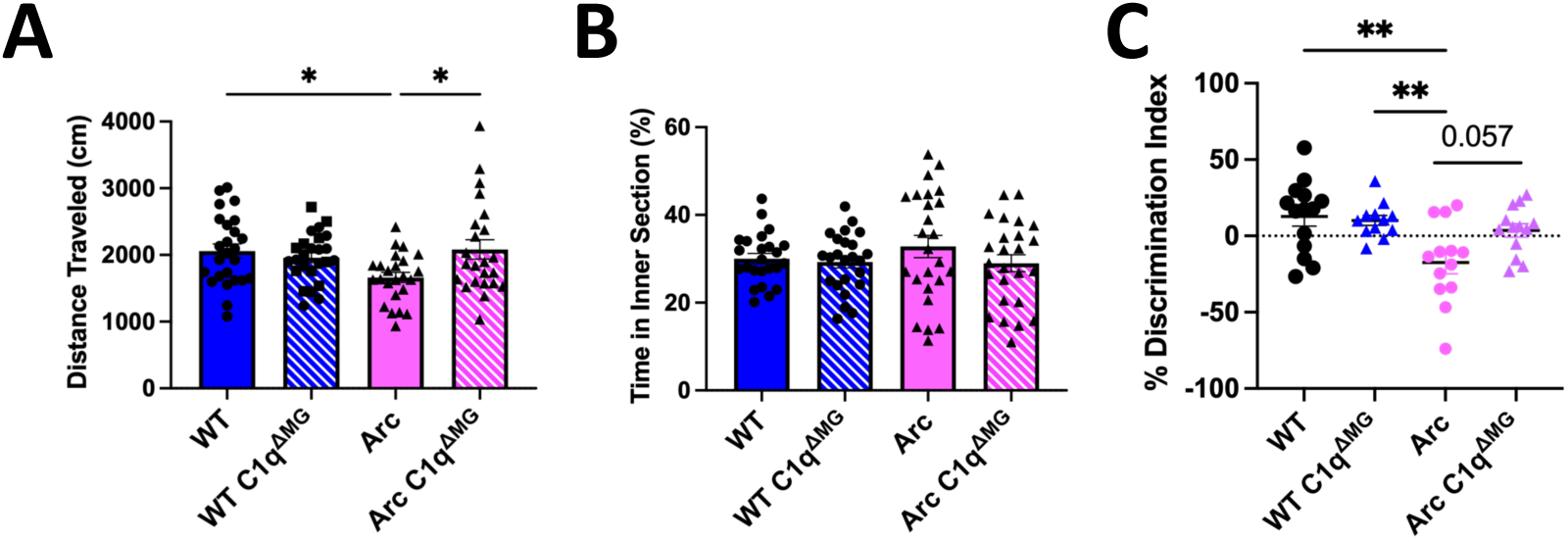
Adult microglial deletion of C1q rescues motor and cognitive impairments in Arctic AD mice. ***A*.** Behavioral tasks revealed a slight deficit in locomotion in 10-month-old Arc mice during 5-minute open field testing (day 1 of habituation to arenas) that was rescued by adult C1q microglial deletion. ***B*.** No changes in anxiety, as measured by the percentage of time spent in the center of the open-field arena were observed between any groups. ***C***. Object location memory testing demonstrated a significant impairment in Arc mice compared to WT controls. Deletion of microglial C1q by 8 weeks of age had a near significant (p=0.057) improvement in the cognitive performance of Arc mice at 10-months of age. Each data point represents 1 animal. Data is expressed as Mean ± SEM; *n* = 22-24 per genotype for open field testing and *n* = 13-15 for object location memory. **p<0.01, ***p<0.001 by One-Way ANOVA followed by Tukey’s post-hoc test.

### Amyloid pathology nor plasma NfL is affected by young adult microglial deletion of C1q

To determine the influence of microglial C1q deficiency on amyloid pathology, we first assessed amyloid plaque burden within the hippocampus with Thioflavin S (ThioS), which binds to the beta-sheet structure of fibrillar Aβ (fAβ) plaques. No difference was observed in fAβ load (ThioS Field Area %) in Arc C1q^ΔMG^ when compared to Arc mice (**Fig 2A-B**) at 10 mo of age. Despite no change in overall amyloid burden or number (**Fig 2C**), Arc C1q^ΔMG^ mice displayed a significance (21%) increase in the average overall size of amyloid plaques (**Fig 2D**). Total hippocampal Aβ40 and Aβ42 was measured in detergent soluble and insoluble fractions. In concordance with overall hippocampal plaque burden and number, no differences were observed in either soluble (**Fig 2E-F**) or insoluble (**Fig 2G-H**) amyloid peptides between Arc C1q^ΔMG^ and Arc mice. These results indicate that while microglial C1q may contribute to the compaction of fibrillar amyloid, it does not significantly influence amyloidosis in the Arc mouse model (Boyett et al., 2003).

**Figure 2:**
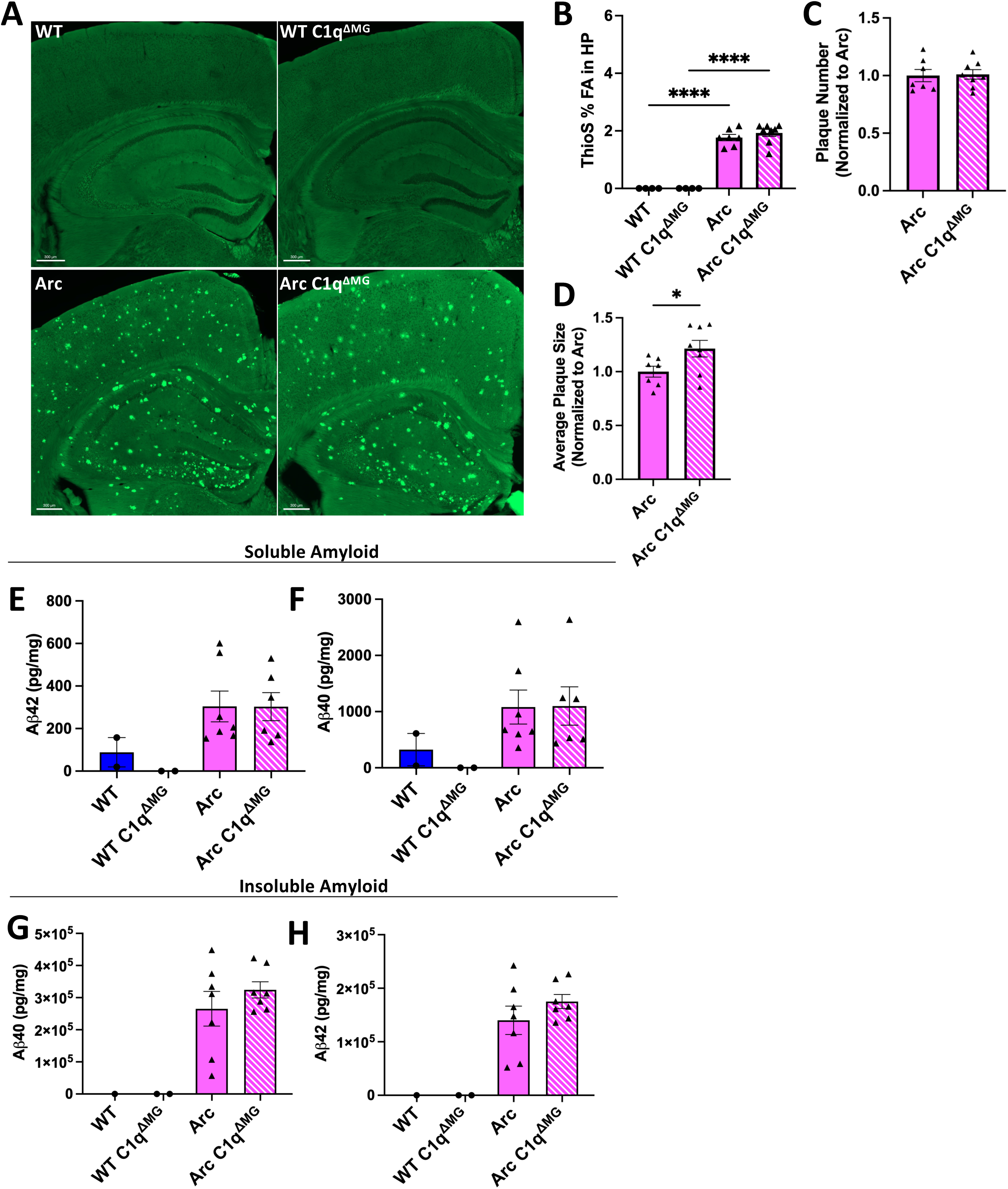
Young adult microglial deletion of C1q fails to alter hippocampal amyloid burden. ***A.*** Representative images of amyloid plaques (Thioflavin S; green) in the hippocampus and cortex. Scale bar: 300μm. **B-D.** Imaris quantification of Thioflavin S field area (***B***), number of plaques (***C***), and average plaque size (***D***) in the hippocampus of WT and Arc mice with and without microglial C1q deletion. Each data point represents the average of 2 sections, *n* = 4 - 8 mice/genotype and is expressed as Mean ± SEM. *p<0.05 by student’s t-test. **E-H.** Levels of soluble and insoluble Aβ were quantified in dissected hippocampi via MesoScale Multiplex technology. No differences in soluble (***E-F***) and insoluble (***G-H***) Aβ40 and Aβ42 were found in Arc mice following microglial deletion of C1q. Each data point represents 1 animal. Data are presented as Mean ± SEM; *n* = 1 –2 for WT and *n* = 6 – 7 for Arc genotypes. ****p<0.0001 by One-Way ANOVA followed by Tukey’s post-hoc test.

Plasma NfL is a common biomarker to assess overall neurodegeneration (Giacomucci et al., 2022). Furthermore, plasma C1q and NfL have been correlated to disease severity in both Huntington’s disease (Tassoni et al., 2022) and frontotemporal dementia (van der Ende et al., 2022) and outcome in traumatic brain injury (Butler et al., 2025). Here, plasma NfL levels at 7 and 10 months of age were quantified to determine if the young adult deletion of microglial C1q influenced overall neurodegeneration. When assessed at 7m of age, Arc mice had a trending increase in plasma NfL compared to WT mice (p=0.06; **Fig S2**) while plasma NfL levels were not elevated in Arc C1q^ΔMG^ compared to WT C1q^ΔMG^ mice. However, by 10m of age both Arc and Arc C1q^ΔMG^ animals had elevated plasma NfL compared to their respective controls (**Fig S2)**. This suggests that while microglial C1q deletion may temporarily suppress an increase in plasma NfL, it does not lead to a sustained reduction.

### Young microglial deletion of C1q reduces levels of C3 in Arc brain, while microglia Iba1, astrocyte GFAP and C5aR1 expression remained unaffected

C3 is known to be elevated in AD patients and AD mouse models (Shi et al., 2015; Wu et al., 2019; Du et al., 2025), including in Arc mice (Carvalho et al., 2022; Schartz et al., 2024). C3 is primarily synthesized by astrocytes in response to inflammatory mediators secreted by microglia, with C1q being identified as a potential key inducer of astrocytic C3 production and induction of neurotoxic astrocytes (Liddelow et al., 2017). To investigate changes in C3 expression and astrocyte reactivity following microglial deletion of C1q, brain sections were stained for C3, GFAP, and Amylo-Glo. Quantitative analysis of the hippocampus demonstrated Arc mice had a significant increase in C3 and GFAP expression compared to WT mice (**Fig S3A)** but reduced C3 reactivity in Arc C1q^ΔMG^ compared to Arc mice (38%; p<0.01; **Fig 3A-B**;). Despite this significant reduction in C3 levels, only a trending reduction in GFAP level was observed (10%; p=0.08; **Fig 3C**). Together, this data suggests the loss of microglial C1q alters the polarization of disease associated astrocytes.

**Figure 3.**
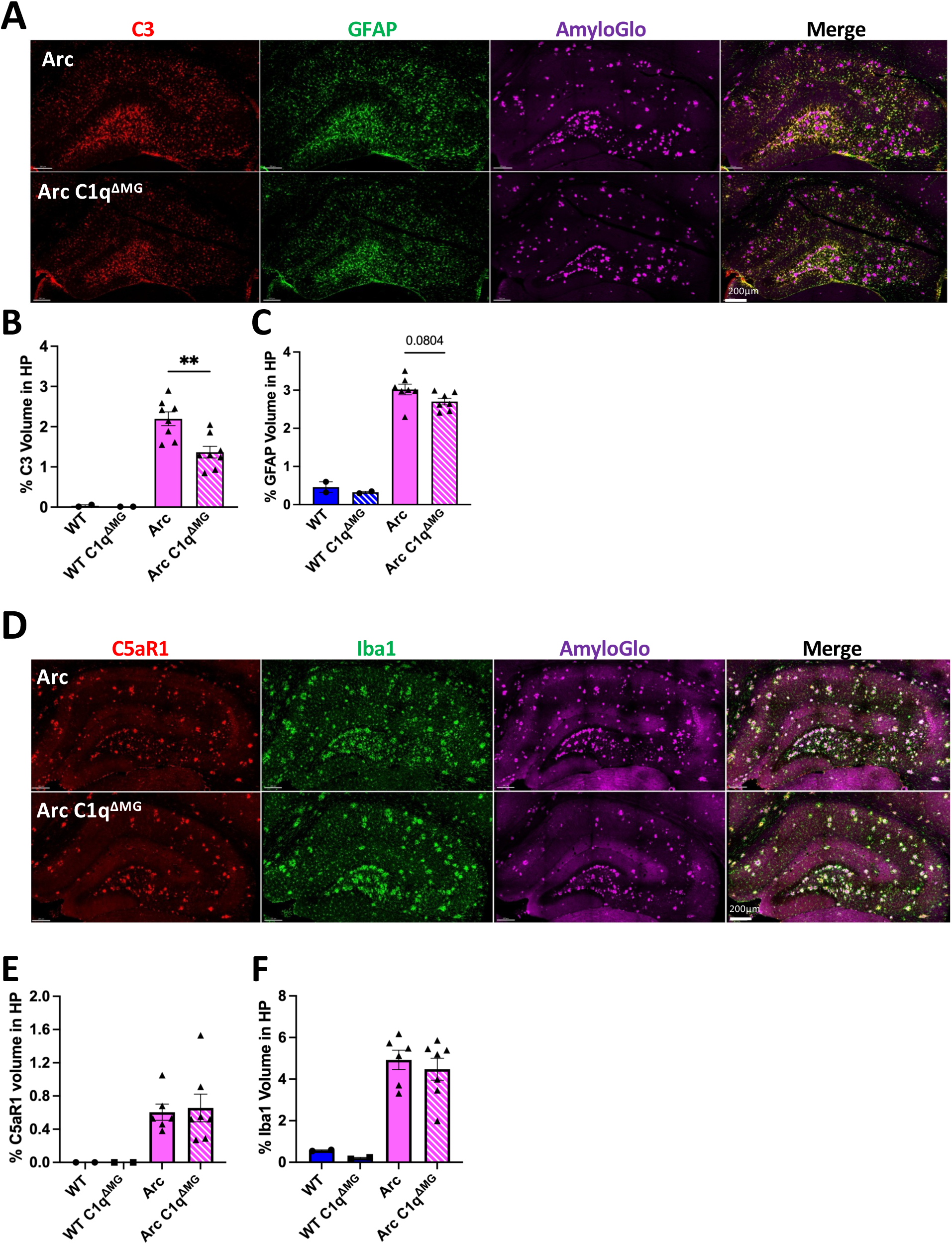
Young microglial deletion of C1q significantly reduces C3 expression but fails to alter glial reactivity or expression of C5aR1. ***A.*** Representative whole hippocampal z stacks of C3 (red), GFAP (green), and AmyloGlo (magenta), and merged images at 20X magnification. Quantification of percent volume of hippocampus of C3 (***B***), or GFAP (***C***). ***D.*** Representative whole hippocampal z stacks of C5aR1 (red), Iba1 (green), and AmyloGlo (magenta), and merged images at 20X magnification. Quantification of percent volume of hippocampus of C5aR1 (***E***), or Iba1 (***F***). Scale bar represents 200μm for all images. T-test between Arc – Arc C1q^ΔMG^ for IF experiments. Data are shown Mean ± SEM of the average of 2 sections, with 2 mice for WT genotypes and 6-8 mice for Arc genotypes. **p<0.01.

Microglia are the primary producers of C1q (Fonseca et al., 2017) and the primary expressors of the complement receptor C5aR1 in AD (Hernandez et al., 2017; Schartz et al., 2024). When activated by its ligand, C5a, C5aR1 activation in microglia promotes inflammatory signaling and chemotaxis of additional immune cells that contributes to AD progression (reviewed in Zelek and Tenner (2025)). To assess the impact of young adult microglial C1q deletion on microglial reactivity and C5aR1 expression, we quantified C5aR1 and Iba1 expression in the hippocampus at 10-months of age. As expected, both C5aR1 and Iba1 expression were increased in Arc mice compared to WT mice (**Fig S2B**). In Arc mice, microglial C1q deletion did not lead to reductions in either C5aR1 or Iba1 immunoreactivity (**Fig 3D-F**). These results support previous reports that microglial C5aR1 upregulation occurs as an early response to injury (Carvalho et al., 2022). These data also indicate that microglial C1q is not a major contributor to the hypertrophic morphological state of microglia.

### Microglial deletion of C1q does not reduce transcription of select complement genes in the Arctic hippocampus

C4, another component of the classical complement cascade, is induced in the brain in response to injury (Tenner and Petrisko, 2025) and its cleavage at the protein level is required to form the classical pathway C3 convertase which participates in synaptic pruning. To determine if reduced C3 gene expression was occurring at the transcriptional level and to determine if loss of microglial C1q influenced expression of C3, C4, or C5aR1. qPCR for these components were performed. Despite the decrease in C3 protein reactivity in the hippocampus (**Fig 3C**), Arc C1q^ΔMG^ hippocampi had similar C3 gene expression to Arc mice (**Fig 4A)** at 10 months of age, suggesting that the decrease in C3 expression is occurring at the translational or post-translational stage. C4 mRNA expression remained elevated in both the Arc and Arc C1q^ΔMG^ hippocampus (**Fig 4B**) relative to wild type, suggesting its transcription is regulated independently of microglial C1q. Finally, matching our protein expression data (**Fig 3E**), C5aR1 gene expression is strongly induced in Arc mice but no difference was observed between Arc and Arc C1q^ΔMG^ hippocampi (**Fig 4C**) at 10 months of age.

**Figure 4.**
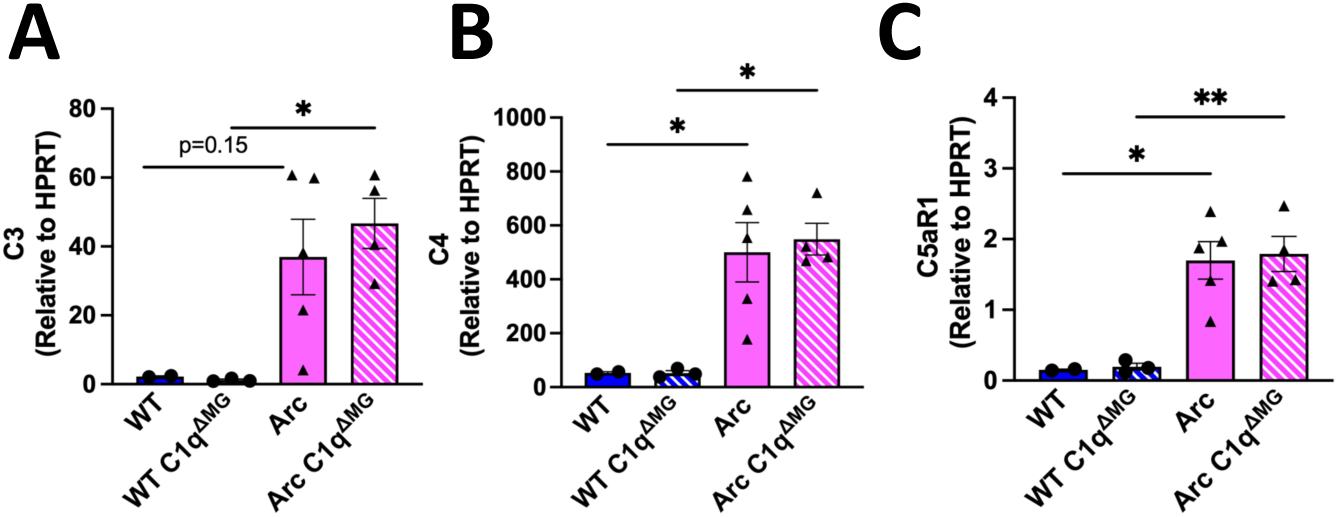
Hippocampal gene expression of C3, C4, and C5aR1 are not altered by loss of microglial C1q. Gene expression of *C3*(***A***), *C4* (***B***), and *C5aR1* (***C***) normalized to *HPRT* expression at 10 mo of age. Each data point represents a single animal, ran in triplicate. *p<0.05, **p<0.01. One-way ANOVA followed by Tukey’s post-hoc test.

### Engulfment of synapses is reduced in Arctic AD mice following young adult microglial deletion of C1q

C1q has a well-established role in the tagging of synapses for engulfment by microglia both in health and disease (Schafer et al., 2012; Hong et al., 2016; Gyorffy et al., 2018; Kovacs et al., 2020; Gomez-Arboledas et al., 2024). Here, the impact of microglial C1q on the engulfment of excitatory synapses (Vglut1) by microglia in the CA1 and CA3 subregions of the hippocampus was assessed. These regions were selected as the CA1-SR and CA3-SL are known to be a part of the tri synaptic circuit, which sequentially connects the entorhinal cortex, dentate gyrus, CA3, and CA1 (Basu and Siegelbaum, 2015).

In the CA1, both Arc and Arc C1q^ΔMG^ mice had increased engagement of Vglut1 excitatory synapses by microglia (Vglut1-Iba1 colocalization) compared to their respective WT controls (p<0.01 WT-Arc; p<0.05 WT C1q^ΔMG^ -Arc C1q^ΔMG^; **Fig 5A-B**). In the CA3 subregion, Arc mice again showed increased engagement of Vglut1 by microglia (Vglut1-Iba1 colocalization) compared to WT mice (**Fig 5D-E**; p<0.0001) at 10 mo of age. While elevated in the CA3, Vglut1-Iba1 colocalization of Arc mice lacking C1q did not significantly differ from WT C1q^ΔMG^ (**Fig 5E**; p=0.17). No differences in Vglut1-Iba1 colocalization were observed in either the CA1 or CA3 between Arc C1q^ΔMG^ and Arc animals, suggesting that microglial C1q is not required for the physical interaction between microglia and Vglut1+ synapses.

**Figure 5:**
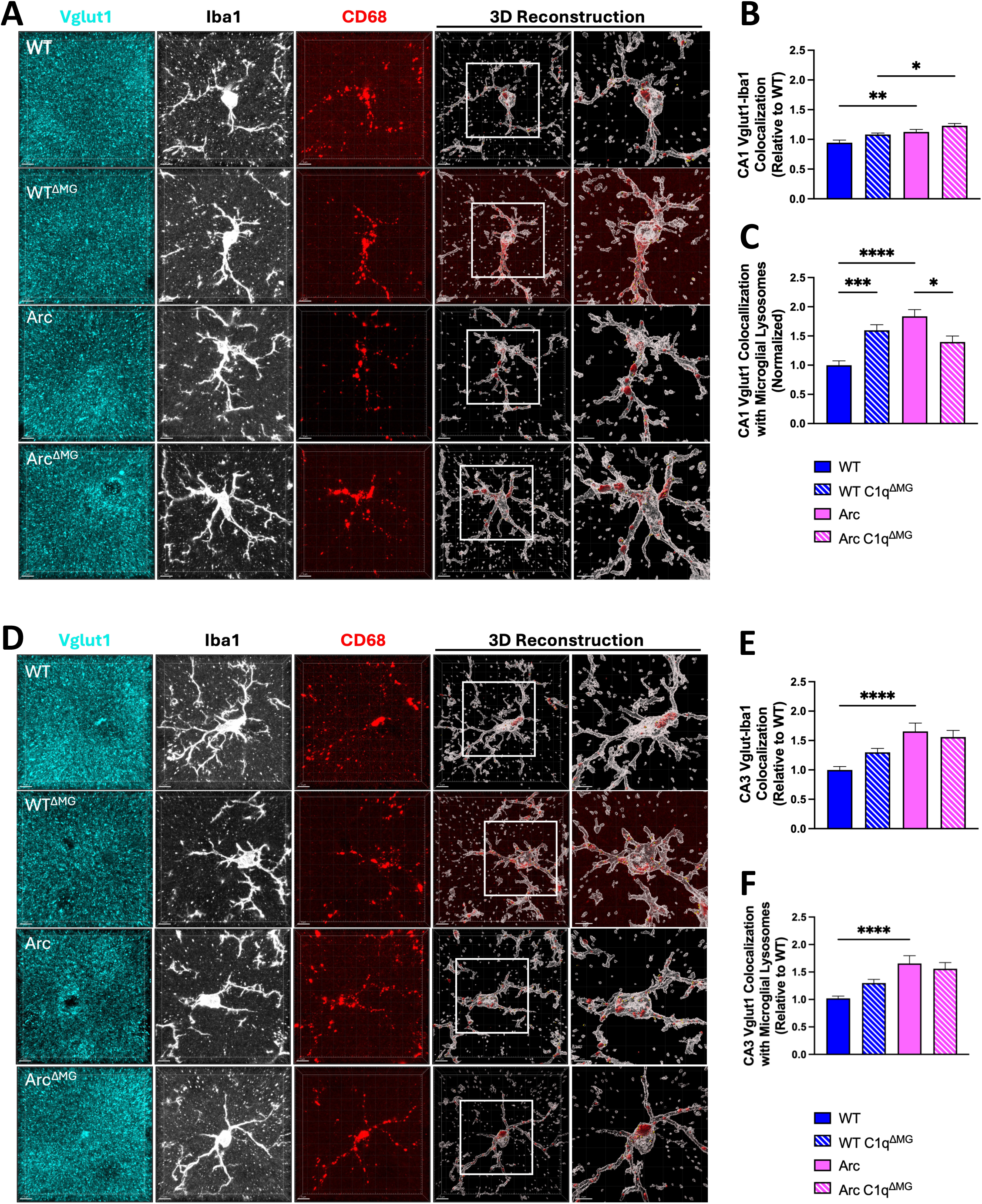
Young microglial deletion of C1q reduced Vglut1 synaptic engulfment in the CA1 but not CA3 subregions of the hippocampus. ***A.*** Representative confocal images and 3D reconstruction with surface rendering using Imaris of individual CA1 microglia cells (Iba1+; white), lysosomes (CD68; red), and presynaptic (Vglut1+; cyan) spots. ***B.*** Quantitative analysis of Vglut1+ presynaptic puncta colocalized with microglia in the CA1 region. ***C.*** Quantitative analysis of Vglut1+ presynaptic puncta localized within microglial lysosomes in the CA1. ***D.*** Representative confocal images and 3D reconstruction with surface rendering using Imaris of individual CA3 microglia cells (Iba1+; white), lysosomes (CD68; red), and presynaptic (Vglut1+; cyan) spots. ***E.*** Quantitative analysis of Vglut1+ presynaptic puncta colocalized with microglia in the CA3 region ***F.*** Quantitative analysis of Vglut1+ presynaptic puncta localized within microglial lysosomes in the CA3 region. Scale bar: 5μm, 4μm insert. Data are shown as Mean ± SEM of 12 individual microglia cells/mouse per region and *n = 4 - 7* per genotype. *p<0.05, **p<0.01, ***p<0.001, ****p<0.0001 by One-Way ANOVA followed by Tukey’s post-hoc test.

As anticipated, Arc mice demonstrated increased Vglut1 within microglial lysosomes compared to WT mice in both CA1 (**Fig 5C**; p<0.0001) and CA3 regions (**Fig 5F**; p<0.0001). However, the loss of C1q led to a significant reduction in the amount of Vglut1 present within microglial lysosomes (Vglut1-Iba1-CD68) in Arc C1q^ΔMG^ mice compared to normal Arc mice (p<0.05) in the CA1 but not CA3 region. Constitutive C1q knockout mice have been shown to have enhanced synaptic connectivity compared to WT mice (Ma et al., 2013); surprisingly, WT C1q^ΔMG^ mice demonstrated an increase in Vglut1 within microglial lysosomes compared to WT mice (p<0.001) in the CA1, which may be the result of altered clearance efficiency.

Together, this data suggests that excessive microglial synaptic pruning seen at the late (10m) stage of AD can be partially rescued by young adult microglial deletion of C1q but in a region dependent manner.

### Young adult deletion of C1q in microglia reduces microglial response to plaques, resulting in reduced phagocytosis of amyloid by microglia

Microglia are known to both aid in the compaction of fAβ and preferentially engulf diffuse Aβ plaques (Condello et al., 2018). To investigate if microglial C1q deletion would alter phagocytosis of amyloid, the engagement of microglia with amyloid and the amount of Aβ within microglial CD68+ lysosomes surrounding plaques were quantified throughout the Arc and Arc C1q^ΔMG^ hippocampus. Arc C1q^ΔMG^ mice displayed a trending reduction in microglial engagement (Iba1-6E10 colocalization) of amyloid compared to Arc mice (p=0.07, 17%; **Fig 6A,B**). Furthermore, Arc C1q^ΔMG^ mice displayed a significant 50% decrease in the amount of amyloid localized within microglial lysosomes (Iba1-6E10-CD68 colocalization) compared to Arc mice (p<0.0001, **Fig 6C**). The reduction in amyloid phagocytosis in Arc C1q^ΔMG^ mice aligns with the observed 17% reduction in Iba1 volume (p<0.05; **Fig 6D**) surrounding plaques as well as the 39% reduction in CD68 reactivity (p<0.0001, **Fig. 6E**); however, if the reduction in CD68 is a direct effect of the loss of C1q or a reflection of the reduction in microglial volume surrounding the plaques remains to be determined.

**Figure 6:**
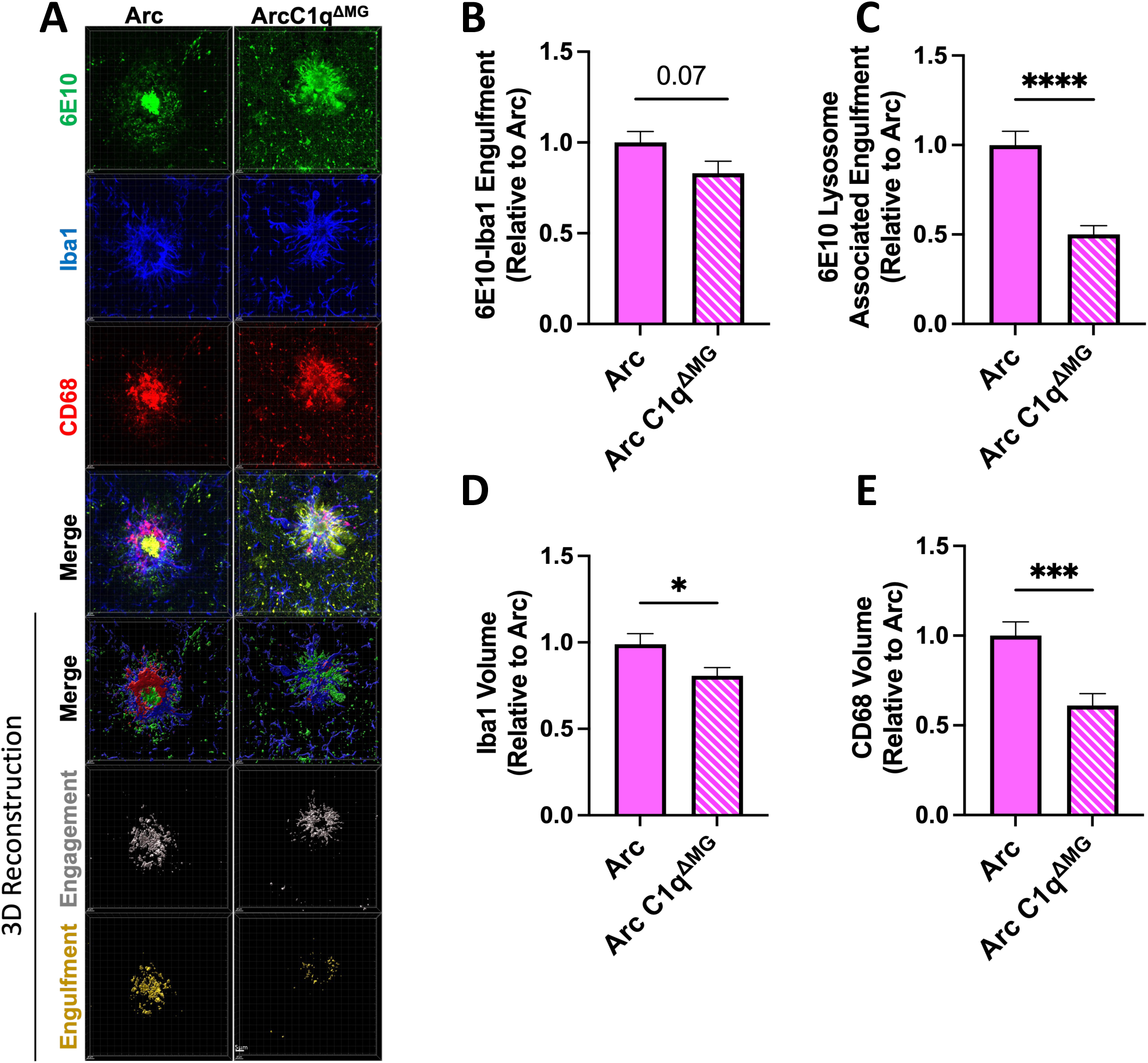
Microglial phagocytosis of amyloid is decreased following young adult microglial deletion of C1q. ***A***. Representative confocal images of amyloid plaques (6E10; green), microglial cells (Iba1; blue), and microglial lysosomes (CD68; red) in the hippocampus of Arc (left) and Arc C1q^ΔMG^ (right) mice. Scale bar: 5μm. ***B.*** Imaris 3D-reconsruction images showed a trend towards reduced internalization of Aβ by microglial cells in Arc C1q^ΔMG^ mice compared to Arc mice (6E10-Iba1 colocalization). ***C.*** 3D Imaris quantification demonstrated reduced Aβ in microglial lysosomes (6E10-Iba1-CD68 colocalization) in Arc C1q^ΔMG^ mice. Microglial deletion of C1q also lead to a reduced volume of microglial cells (***D***) surrounding the amyloid plaques and an overall reduction of CD68 (***E***). Data expressed as Mean ± SEM of 20 hippocampal plaques per mouse, with *n* = 4 – 6 mice per genotype and normalized to Arc genotype. *p<0.05, ***p<0.001, ****p<0.0001 by One-Way ANOVA followed by Tukey’s post-hoc test.

### Young adult deletion of C1q in microglia reduces amyloid phagocytosis by astrocytes but does not alter astrocytic interactions with plaques

In contrast to microglia, astrocytic association with amyloid (GFAP-6E10 colocalization) did not differ between Arc and Arc C1q^ΔMG^ animals (**Fig 7A-B**). Concordant with this data, no change was observed in the overall volume of astrocytes surrounding the plaques (**Fig 7D**). Despite no change in the amount of interaction between astrocytes and amyloid plaques, Arc C1q^ΔMG^ mice displayed a significant reduction (45%) in the amount of amyloid located within astrocytic Lamp2+ lysosomes (p<0.0001; **Fig 7C**) when compared to Arc animals. Corresponding to this result, Arc C1q^ΔMG^ animals had a significant decrease (19%) in the volume of the lysosome marker, Lamp2 (p<0.01; **Fig 7E)**, surrounding amyloid plaques compared to Arc mice. Taken together, this indicates that microglial C1q contributes to the astrocytic phagocytosis of amyloid. The observed reduction in amyloid phagocytosis by both microglia and astrocytes in Arc C1q^ΔMG^ mice despite unchanged hippocampal amyloid burden (**Fig 2B**) may reflect a broader suppression of neuroinflammatory signaling that may drive APP processing as well as microglial engagement with plaques (Cho et al., 2007; Sutinen et al., 2012; Alasmari et al., 2018).

**Figure 7:**
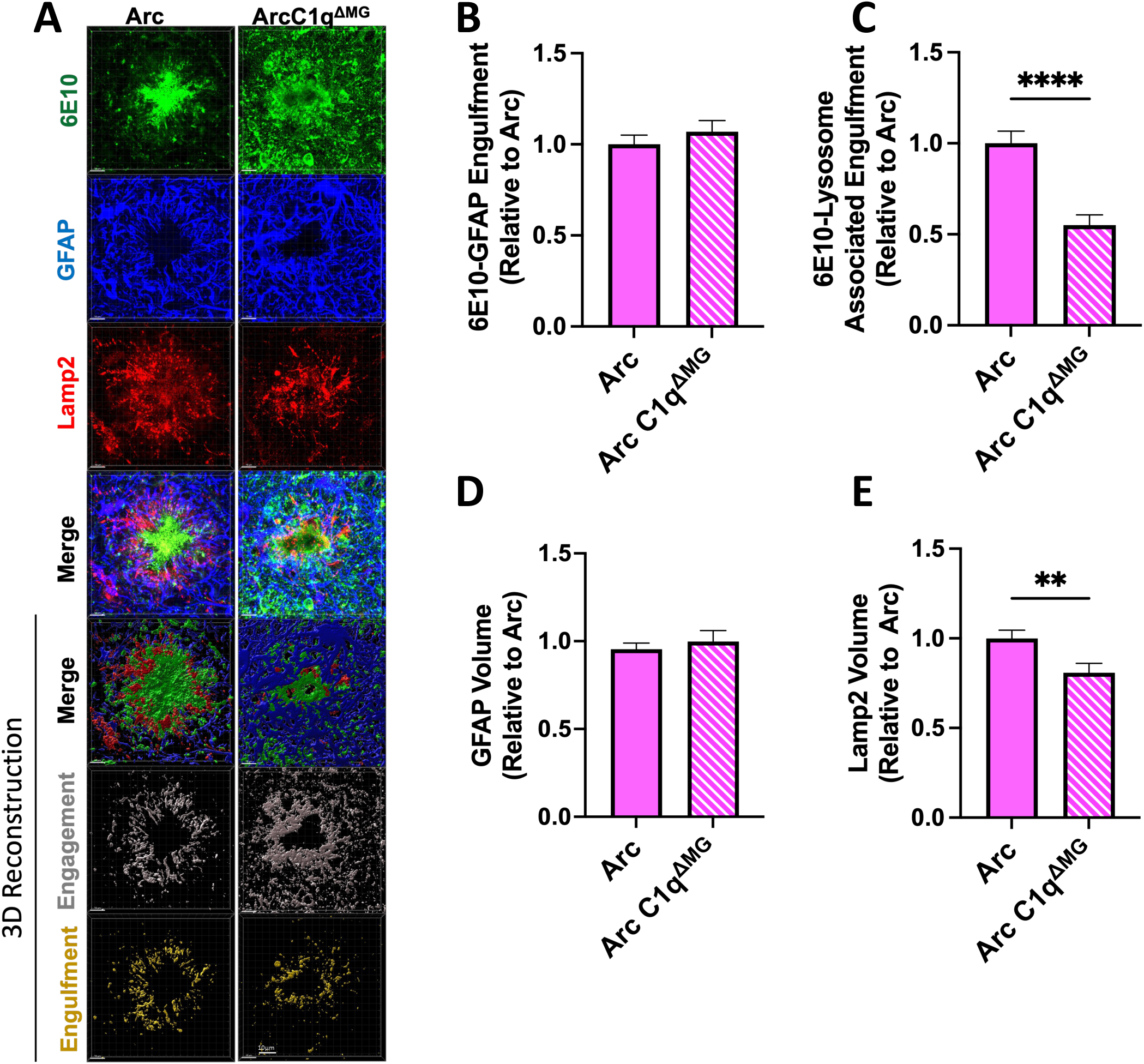
Astrocyte phagocytosis of amyloid is decreased following young adult microglial deletion of C1q. ***A***. Representative confocal images of amyloid plaques (6E10; green), astrocytes cells (GFAP; blue), and astrocytic lysosomes (Lamp2; red) in the hippocampus of Arc (left) and Arc C1q^ΔMG^ (right) mice. Scale bar: 5μm. ***B.*** Imaris 3D-reconsruction images showed no difference in the internalization of Aβ by astrocytes in Arc C1q^ΔMG^ mice compared to Arc mice (6E10-Iba1 colocalization). ***C.*** 3D Imaris quantification demonstrated reduced Aβ in astrocytic lysosomes (6E10-GFAP-Lamp2 colocalization) in Arc C1q^ΔMG^ mice. Microglial deletion of C1q did not alter the volume of astrocytes (***D***) surrounding amyloid plaques but did lead to an overall reduction of Lamp2 (***E***). Data expressed as Mean ± SEM of 20 hippocampal plaques per mouse, with *n* = 4 – 6 mice per genotype and normalized to Arc genotype. **p<0.01, ****p<0.0001 by One-Way ANOVA followed by Tukey’s post-hoc test.

## Discussion

The complement system, particularly the classical pathway initiated by fibrillar amyloid, hyperphosphorylated tau, and damage associated molecular patterns (DAMPs) binding to C1, plays a critical role in AD pathogenesis through its contributions to synaptic pruning and neuroinflammation. C1q expression is upregulated as an early response to injury and activates astrocytes to produce C3, enabling synaptic pruning of “weak” synapses which can lead to cognitive decline. Microglia are the primary producers of C1q in the brain, and understanding the specific role of microglial-derived C1q is crucial for elucidating AD mechanisms and developing targeted therapeutics. The present study demonstrates that young adult microglial deletion of C1q in a mouse model of amyloidosis results in a long-term net benefit, including preserved cognitive function, decreased synaptic pruning, and reduced astrocytic C3 expression despite minimal effects on overall amyloid burden. These results further support the hypothesis that it is the immune response – including activation of a powerful multifaceted complement cascade by amyloid – rather than amyloid itself that is a driver of synaptic loss and cognitive decline in AD (Hernandez et al., 2017; Miao et al., 2023; Ayyubova and Fazal, 2024; Schartz et al., 2024).

Congruent to our results here, a previous study by Fonseca et al. (2004) demonstrated constitutive genetic C1q deletion in both the Tg2576 and APP/PS1 AD mouse models had no impact on amyloid pathology in aged mice. While synaptic engulfment was not directly assessed in that study, the authors observed a preservation of the synaptic marker synaptophysin as well as the cytoskeletal marker Map2 in aged Tg2576 mice lacking C1q. This result correlates with our observed reduction in Vglut1+ within microglial lysosomes within the CA1 as well as preservation of spatial memory at 10 months of age in Arc C1q^ΔMG^ animals. Furthermore, Fonseca et al. (2004) demonstrated that the deletion of C1q significantly reduced astrocyte and microglia activation by ∼50% in both mouse models. In contrast, our young adult microglial C1q deletion in the aggressive Arc model demonstrated no change in either astrocyte GFAP or microglial Iba1 protein expression, reiterating that expression levels of these glial cellular markers alone are not always a reliable surrogate for functional activation states. Indeed, the reduction in astrocytic C3 (**Fig 3B)** and in microglia surrounding plaques (**Fig 6D**) suggests transcriptional reprogramming of Arc C1q^ΔMG^ astrocytes and microglia. Overall, our study demonstrates the importance of assessing both the spatial and temporal role of C1q throughout AD.

Additionally, glial phagolysosome colocalization of amyloid is significantly reduced in Arc C1q^ΔMG^ microglia **(Fig 6**) and astrocytes **(Fig 7**), alongside a reduction in Iba1 volume surrounding fAβ plaques. This may reflect a general modulation of microglia and astrocytes to a less inflammatory state, as supported by our reduced C3 expression, or increased digestion/processing of the amyloid cargo. The dynamic interplay between pro- and anti-inflammatory signaling pathways is known to drive amyloid plaque accumulation (Lee et al., 2008; Chouhan et al., 2021). Alternatively, the C1q enhanced induction of Lrp1b (Benoit et al., 2013), which reduces amyloid internalization/trafficking/processing, may explain why amyloid levels are unchanged at 10 months of age despite reduced lysosomal colocalization in Arc C1q^ΔMG^ animals.

We also observed a 21% increase in plaque size following young adult microglial C1q deletion, which would be compatible with a model in which C1q contributes to amyloid compaction rather than (or in addition to) clearance (**Fig 2D**). This finding aligns with previous reports that C1q can promote the formation of dense-core plaques (Boyett et al., 2003). Diffuse plaques have been shown to be more easily internalized by microglia than dense core plaques; however, it has been suggested that this microglial internalization is a mechanism to sequester and compact amyloid into dense core plaques as a means of limiting the effects of amyloid on neurons and thus the loss of C1q prevented the internalization and conversion of amyloid to a dense core morphology (Huang et al., 2021). Higher levels of diffuse to dense-core amyloid plaques have also been associated with reduced cognitive impairment (Liu et al., 2022). While our results support C1q’s role in microglial compaction of amyloid, the observed improvement in cognition may also be due to the long-term anti-inflammatory effects of C1q deletion.

The molecular mechanisms underlying the observed reduction in amyloid phagocytosis likely occur through different pathways in microglia and astrocytes. The 50% reduction in microglial amyloid phagocytosis may reflect the direct role of C1q as an opsonin as well as indirectly due to the generation of C4b/C3b/iC3b facilitating microglial recognition and engulfment of amyloid deposits through complement receptors (Fu et al., 2012; Lv et al., 2024). In contrast, astrocytes express the phagocytic receptors Megf10 (Multiple EGF-like-domains 10) and MerTK (Mer tyrosine kinase) (Chung et al., 2013). MerTK binds to phosphatidylserine (PtdSer), which can also bind C1q, whereas Megf10, whose expression is increased in response to injury, can bind C1q directly. Acting together, MerTK and Megf10 signaling promote astrocytic engulfment of synapses (Chung et al., 2013; Iram et al., 2016; Shi et al., 2021; Zhuang et al., 2023).Therefore, the 45% reduction in amyloid phagocytosis by astrocytes may be due to disrupted C1q-Megf10 - MerTK signaling pathways and the general disruption of microglial-astrocyte communication following C1q deletion may explain why both cell types showed impaired clearance function and reduced expression of lysosomal makers in plaque associated glia. The fact that reduced colocalization of amyloid with CD68 or Lamp2 is nevertheless associated with improved cognition, again supports the hypothesis that other downstream complement activation products (such as C5a and C5b-9) that would be reduced in the absence of C1q are indeed the more proximal factors leading to neuroinflammation induced cognitive dysfunction (Schartz et al., 2024; Zelek et al., 2025).

We observed region-specific effects of early microglial C1q deletion on microglial synaptic engulfment. Specifically, young adult microglial deletion reduced the amount of Vglut1 inside microglial lysosomes within the CA1 region(**Fig 5C**), while having minimal impact in the CA3 region (**Fig 5F**). This differential response may reflect the distinct anatomical and functional properties of these hippocampal subregions within the trisynaptic circuit (Basu and Siegelbaum, 2015). Alternatively, in the Arc model, hippocampal amyloid depositions begin around 3 months of age and typically initiates in the dentate gyrus. Given the direct projections from the dentate to CA3, the CA3 may be more vulnerable to early AD pathology, such that C1q deletion here is not early enough to fully mitigate increased synaptic engulfment.

Our observation that microglial C1q deletion led to a 38% reduction in hippocampal C3 levels (**Fig 3B**) supports previous observations of C1q as an inducer of astrocytic C3 production (Liddelow et al., 2017; Guttikonda et al., 2021). Although the signaling mechanisms driving C1q-induced astrocytic C3 production are not fully defined, several studies have demonstrated that astrocytic production of C3 (and subsequent cleavage into C3a) is critical for mediating the interactions between astrocytes and microglial via C3a-C3aR signaling (Chen et al., 2020; Wei et al., 2021). These findings collectively underscore the intricate bidirectional communication between astrocytes and microglia and position C1q-C3a-C3aR signaling as a key regulatory pathway in glial cell crosstalk and neuroinflammatory responses.

The absence of changes in C5aR1 expression following microglial C1q deletion (**Fig 3E**) is consistent with C5aR1 serving as an early injury response marker that may be regulated independently of C1q signaling (Carvalho et al., 2022; Schartz et al., 2024). However, the absence of C1q reduces the generation of C5a via classical pathway activation thereby preventing pro-inflammatory C5aR1 signaling in microglia and further limiting the inflammatory communication between microglia and astrocytes. Although the overall hippocampal volume of Iba1 remained unchanged following C1q deletion (**Fig 3F**), the observed alterations in both synaptic and amyloid engulfment suggest that microglial gene expression and/or proteome profiles are likely modified despite the preservation of overall microglial morphology and density (Dejanovic et al., 2022; Scott-Hewitt et al., 2024), consistent with results from other complement genetic deletions (Shi et al., 2015; Hernandez et al., 2017; Carvalho et al., 2022).

This work demonstrates the potential of long-term cognitive and likely neuroinflammatory benefits from microglial specific C1q deletion beginning in young adulthood in AD. However, several limitations of this study warrant consideration. The Arc mouse model, while valuable for studying amyloid-related pathology, may not fully recapitulate the complexity of human AD and future studies should utilize newer humanized knock-in mouse models (Saito et al., 2014; Baglietto-Vargas et al., 2021; Xia et al., 2022).

A study by Dejanovic et al. (2022) demonstrated microglia preferentially engulf inhibitory synapses while astrocytes preferentially engulf excitatory synapses in a C1q-dependent manner, although astrocytes could compensate for impaired microglial phagocytosis of inhibitory synapses in Trem2 (Triggering Receptor Expressed on Myeloid Cells 2) deficient mice. Our study only examined microglial synaptic engulfment of the excitatory pre-synaptic protein, Vglut1. Future studies should further investigate the role of microglial C1q deletion on astrocyte polarization and on ingestion of both excitatory and inhibitory synapses.

Additionally, future studies should examine the time course of neuroinflammation and cognitive change following microglial C1q deletion. While reduced astrocytic C3 and reduced microglial engulfment of synapses and improved cognition were observed at 10 months of age, *in vitro* studies have demonstrated that C1q, in the absence of other downstream complement components, as would occur in the earliest stages of disease, is neuroprotective against amyloid-mediated neuronal cell death (Benoit and Tenner, 2011; Benoit et al., 2013). C1q has also been implicated in myelination during development (Yu et al., 2023), as well as promoting repair following spinal cord injuries, indicating that at times, C1q may be beneficial, particularly acutely (Benavente et al., 2020). That some C1q expression can be beneficial is also supported by results from a human clinical trial in which individuals with Guillain Barrè symptoms were treated with the anti-C1q therapeutic ANX005 (Annexon Biosciences). Patients on the higher, 70mg/kg dose of ANX005 failed to have an improved outcome at the end of the study, whereas individuals on the lower 35mg/kg dose showed improved neurological outcomes (Annexon Biosciences, 2024).This correlated with levels of antiC1q persisting into the recovery phase, suggesting that too much C1q inhibition or inhibiting C1q for long periods may be detrimental.

In summary, this work provides important insights into the temporal dynamics of complement-mediated neuroinflammation and highlights the potential of targeting C1q in Alzheimer’s disease. As previous studies have demonstrated a neuroprotective role for C1q and signaling mechanisms independent of the complement cascade, future studies examining the role of microglial C1q during different stages of AD must be performed. If C1q is indeed neuroprotective in the early stages of AD, anti-C1q therapeutics could worsen cognitive performance if given too early in disease progression. Finally, understanding the full network of C1q-responsive receptors across cell types that may mediate C1q functions across disease stages will be essential for developing comprehensive therapeutic strategies targeting C1q.

## Author Contributions

TJP contributed to experimental design, tamoxifen administration, tissue collection, IF experiments and analysis, MSD assays, and wrote the manuscript. SHC contributed to mice breeding and generation, tamoxifen administration, tissue collection and processing IF experiments and analysis. AGA contributed to interpretation of results. BZ performed IF and analysis. AJT contributed to experimental design and manuscript preparation. All authors reviewed, edited, and approved the manuscript.

## Acknowledgements

We thank Dr. Lennart Mucke (Gladstone Institute of Neurological Disease) for the Arctic mice. We also thank Lauren Chung, Andrian Mendoza-Arvilla, and Megan Garcia for their assistance in sectioning the tissue samples and plasma isolation and Karina Chavez for their assistance in conducting the NfL assay.

## Data availability

Data available upon request.

---

**Figure S1.** Confirmation of C1q deletion in microglia at 10 months of age. **A.** Representative images of C1q staining in the dentate gyrus of the dorsal hippocampus. Scale bar represents 100μm. ***B.*** Quantification of C1q intensity in molecular layer of dentate gyrus. Each data point represents the average mean intensity of 2 sections per animal. *n = 4-8* mice per genotype ***C.*** Complete western blot of C1q and corresponding β-actin blot of hippocampal tissue. The *C1qKO* mouse is a 10-month Arc C1q gene trapped mouse (Fonseca et al., 2017) while the *Ref* sample is a 10-month Arc C1qa^FL/FL^ mouse to allow for normalization across multiple blots. ***D.*** Quantification of C1q hippocampal western blot normalized to β-actin. One-way ANOVA.**p<0.01, ****p<0.0001

**Figure S2.** Young adult microglial C1q deletion transiently suppresses plasma NfL. Plasma NfL was assessed at 7 and 10m of age. Results expressed as pg/mL plasma. *n* = 6-14 per genotype per timepoint with samples run in duplicate. Two-way ANOVA followed by Tukey’s post-hoc test comparing within age only for NfL. *p<0.05, **p<0.01, ***p<0.001

**Figure S3.** Representative images of WT C3/GFAP and C5aR1/Iba1 hippocampal immunohistochemistry. ***A.*** Representative whole hippocampal z stacks of C3 (red), GFAP (green), and AmyloGlo (magenta), and merged images at 20X magnification. ***B.*** Representative whole hippocampal z stacks of C5aR1 (red), Iba1 (green), and AmyloGlo (magenta), and merged images at 20X magnification. Scale bar represents 200μm. Quantification in Figure 1B,C,E,F.

