## Supplementary figures and images for "Young adult microglial deletion of C1q reduces engulfment of synapses and prevents cognitive impairment in an aggressive Alzheimer’s disease mouse model"

### Supplemental Figures

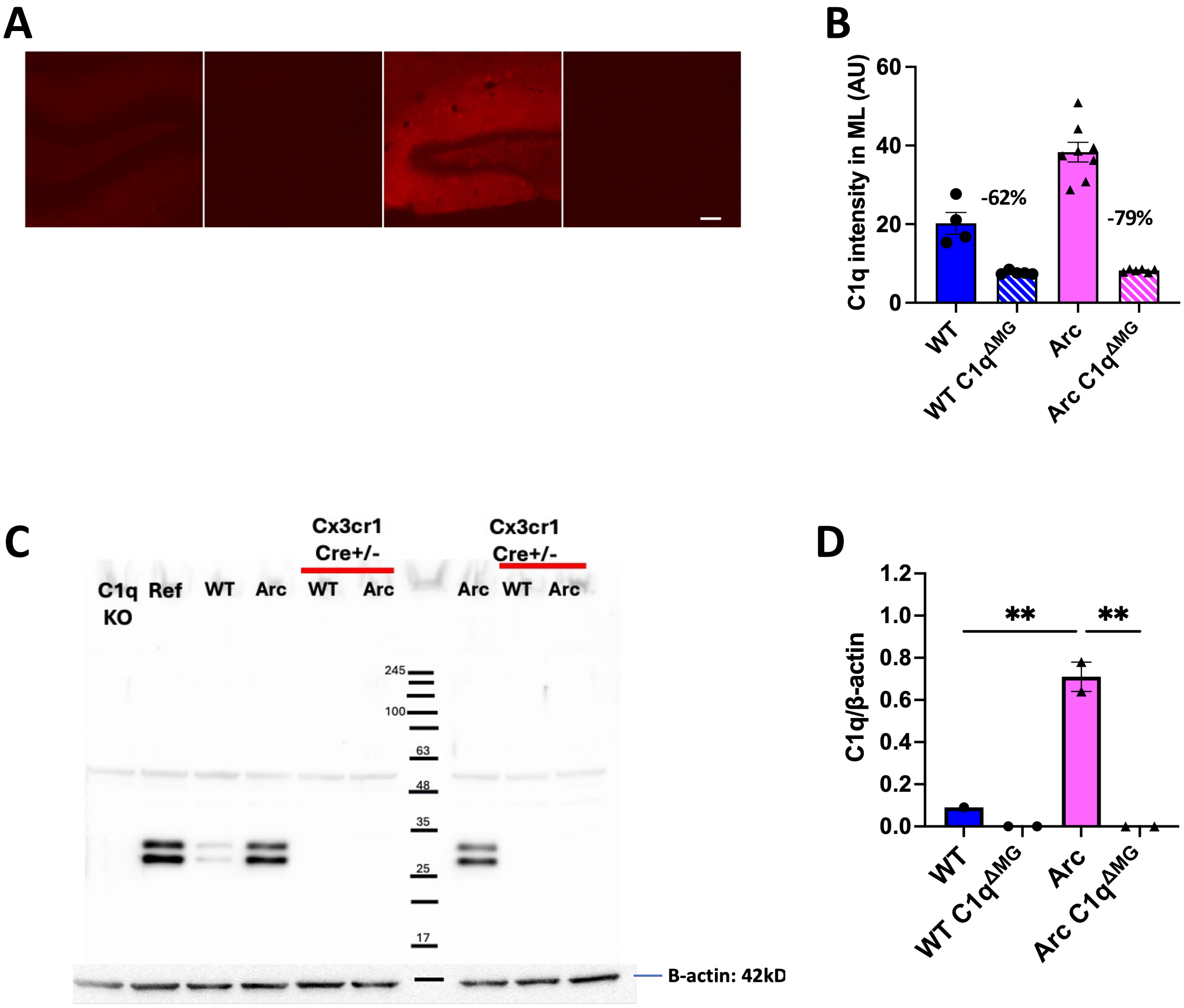

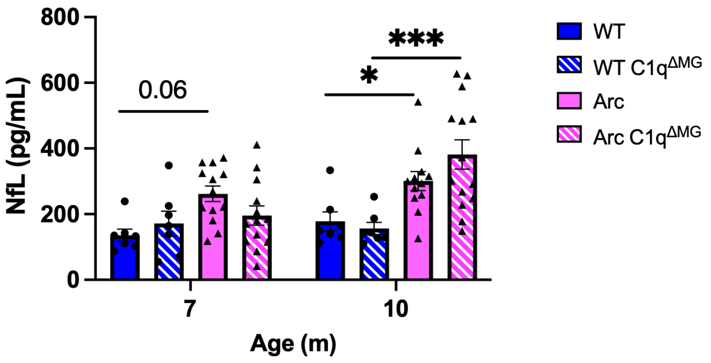

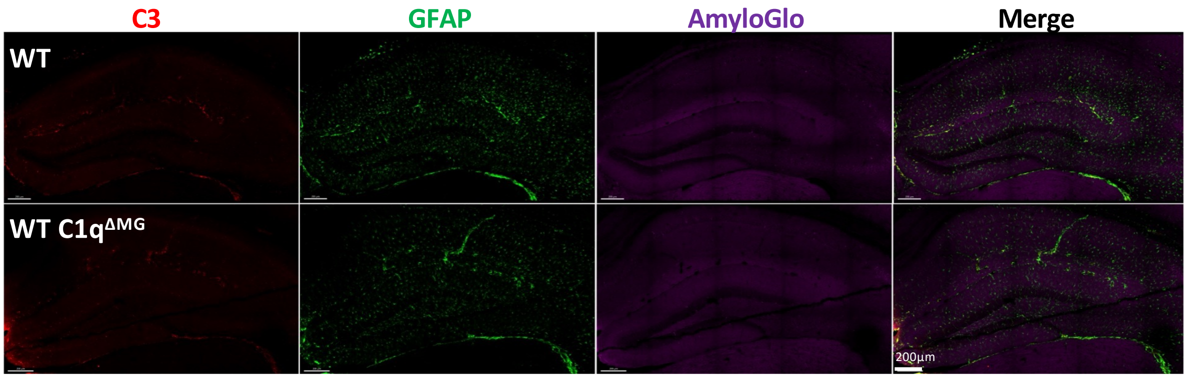

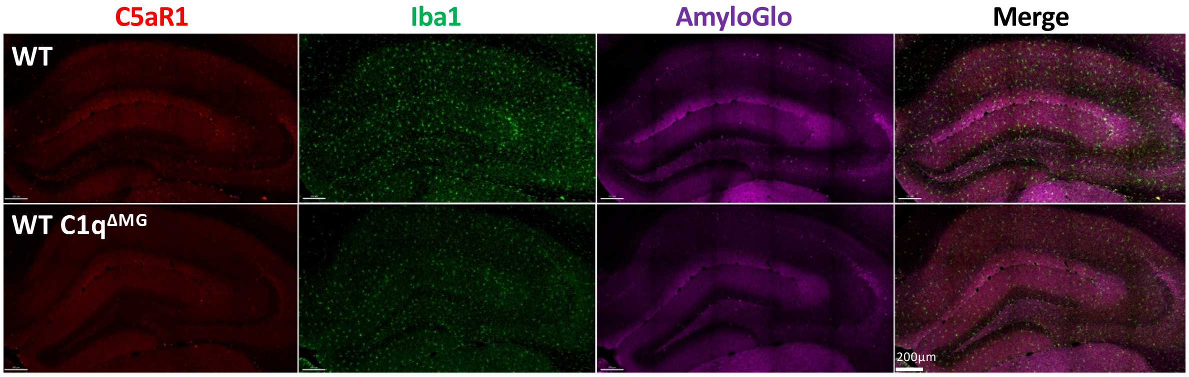
